# A physiology-based model of bile acid metabolism shows altered tissue concentrations after drug administration and in specific genotype subgroups

**DOI:** 10.1101/450965

**Authors:** V. Baier, H. Cordes, C. Thiel, U. Neumann, L.M. Blank, L. Kuepfer

## Abstract

Drug-induced liver injuries (DILI) are an important issue in drug development and patient safety and often lead to termination of drug-development programs or late withdrawals of drugs. Since DILI events are hard to diagnose in preclinical settings, a need for alternative prediction methods such as computational modeling emerges. Impairment of bile acid (BA) metabolism, known as cholestasis, is a frequent form of DILI. Being rather a systemic then a single organ related disease, whole-body physiology-based modeling is a predestined approach for cholestasis modeling. The objectives of the presented study were 1) the development of a physiology-based model for human bile acid metabolism, 2) model validation and characterization for a virtual population, and 3) prediction and quantification of the effects of genetic predispositions and drug interaction on bile acid metabolism. The developed physiology-based bile acid (PBBA) model is based on the standard PBPK model of PKSim® and describes the bile acid circulation in a healthy reference individual. Active processes such as the hepatic synthesis, gallbladder emptying upon meal intake, transition through the gastrointestinal tract, reabsorption into the liver, distribution within the body, and excretion are included. The kinetics of active processes for the surrogate BA glycochenodeoxycholic acid were fitted to time-concentration profiles of blood BA levels reported in literature. The robustness of our PBBA model is underlined by the comparison of simulated plasma BA concentrations in a virtual population of 1,000 healthy individuals with reported data. In addition to plasma concentrations, the PBBA model allows simulations of BA exposure in relevant tissues like the liver and can therefore enhance the mechanistic understanding of cholestasis. This feature was used to analyse the reported increased risk of cholestatic DILI in Benign Recurrent Intrahepatic Cholestasis type 2 (BRIC2) patients. Simulations of the PBBA model suggest a higher susceptibility of BRIC2 patients towards cholestatic DILI due to BA accumulation in hepatocytes. Apart from these intrinsic effects, drug-interactions and their effect on the systemic bile acid metabolism were simulated by combining the PBBA model with a drug PBPK model of cyclosporine A (CsA). The results of which confirmed the reported higher risk of developing DILI as a consequence of CsA intake. Altogether, the presented model enhances our mechanistic understanding of cholestasis, allows the identification of drug-interactions leading to altered BA levels in blood and organs, and could be used to prevent clinical cases of cholestasis and enhance patient safety.

## Introduction

Drug-induced liver injuries (DILI) place a huge burden on health care systems worldwide. About 2-19 incidences per 100,000 habitants occur annually in Europe with symptoms ranging from mild forms such as slightly elevated blood levels of liver enzymes to fatal clinical incidents resulting in acute liver failure (Björnsson 2016; Kaplowitz 2005). Due to this enormous medical relevance, the detection of DILI at an early stage would be highly beneficial, both for a duly termination of treatment with the DILI-causing compound as well as for an early start of therapeutic interventions with curative counteragents. Manifestations of DILI can be differentiated in hepatocellular DILI, cholestatic DILI where cellular damage of the hepatocytes or of the cholangiocytes is the predominant injury, respectively, or a mixture of both (Hamilton et al. 2016). For the categorisation of DILI, current regulatory diagnosis guidelines assess the relative increase in blood plasma levels of the enzymes alanine transferase (ALT) and alkaline phosphatase (ALP). Elevated ALT levels are a general surrogate marker for liver injuries since ALT is released into the blood from the cytoplasm of lysed hepatic cells (Chalasani et al. 2014). In contrast, increased ALP levels are a specific marker for cholestasis since ALP is a degradation product from damaged cholangiocytes in bile ducts (Vinken 2013). It should be noted that both increased ALT and increased ALP levels are merely secondary effects which only become diagnosable once the actual damage has occurred.

Clearly, more indicative markers would be desirable to support an earlier diagnosis of DILI. This, however, requires a mechanistic understanding of the underlying physiological alterations. For hepatocellular DILI a number of in vitro assays as well as computational models are available which allow the analyses of drug-induced responses of intracellular pathways (Kullak-Ublick et al. 2017). Cholestatic DILI is more complex to investigate since it originates from an impaired interplay of multiple organs at the whole-body level, in particular the impaired bile acid (BA) formation and circulation. BAs are endogenous molecules with various functions. With their detergent effects and micelle forming, BAs solubilise excess cholesterol in bile or dietary fat in the intestine and enhance cholesterol excretion and absorption of fat and fat-soluble vitamins. (Jansen et al. 2017) BAs are the driving force of bile formation, as such providing an excretory way for lipophilic substances like bilirubin, steroids, or xenobiotics and their metabolites (Hofmann 1999). In addition, BAs function as signalling molecules in different pathways like homoeostasis of cholesterol, energy and glucose (Houten et al. 2006).

The total BA pool is characterized by a broad spectrum of endogenous BA species. They are either synthesized de novo in liver cells and transformed to corresponding conjugates there or they are actively taken up from hepatic sinusoids following prior enterohepatic. From the liver, BAs are secreted to bile ducts and gallbladder and are further on transported into the gastro-intestinal tract and metabolised by the microbial gut flora. While from the intestinal lumen, they are absorbed into the portal vein and enter the vascular circulation. Enterohepatic circulation of bile acids is hence a systemic process which involves multiple tissues and active transport processes. Impairment of canalicular transporters, such as multidrug resistance protein 1/3, multidrug resistance-associated protein 2 and bile salt export pump (BSEP), results in the potentially toxic accumulation of BAs in liver cells or other tissues (Castro and Pereira Rodrigues 2017; Jackson et al. 2016; Wagner and Trauner 2016).

Such an accumulation of BAs can lead to the clinical symptoms of cholestasis including pruritus and jaundice. Hence, from a medical perspective, BAs would be an ideal biomarker for cholestasis as their blood plasma levels may be increased before the actual cellular damage has occurred. Recent improvements in analytical methods would well enable such a fine-grained analysis of different BA species at the molecular scale for a differential diagnosis of DILI. In clinical practice, these analytics are, however, still not applied as a standard methodology. Since bile acid composition is dependent on the sampling site and plasma profiles may not be representative for specific tissues the identification of bile acid marker profiles for cholestasis is difficult (Eggink et al., 2018). Furthermore, bile acid circulation cannot be described appropriately with any in vitro assay. For this, an assay would require the preservation of not only liver-specific morphology and functionality in an organotypic microenvironment as e.g. in sandwich or spheroidal microtissue cultures (Bell et al. 2016), but also the interplay of further enterohepatic organs. Therefore, even advanced assays can only focus on limited aspects of cholestasis, like the BA passage through parenchymal cells and potential drug interaction with hepatic transporters.

Computational modelling bears the promise to contribute to a mechanistic understanding of the interplay of the various physiological processes underlying the BA metabolism. Such mechanistic models, which would ideally be both knowledge-driven and physiology-based, need to describe enterohepatic circulation of BAs as well as their accumulation in different tissues. Moreover, computational models may be used to simulate physiological concentration profiles that are not accessible experimentally in vivo such as the intracellular space of different organs. In addition, computational models may help to summarise the existing knowledge in a mathematical representation to identify gaps in the current systemic understanding. Likewise, they may be used to identify targeted screening biomarkers for an early indication of specifically cholestatic pathophysiological alterations.

In this study, we present a physiology-based model of bile acid metabolism at the whole-body level based on physiology-based pharmacokinetic (PBPK) modelling. Our model describes the systemic distribution and enterohepatic circulation of glycochenodeoxycholic acid (GCDCA) as a surrogate BA including its synthesis, transport, distribution, and excretion. The developed physiology-based bile acid (PBBA) model was validated with measurements of BA plasma levels from healthy individuals and applied to predict drug interactions and assess genetic predisposition for cholestasis of patients as such representing typical clinical manifestations for DILI and even cases of idiosyncratic cholestasis. In particular, the model was used to analyse shifts in BA levels relative to the reference state of healthy individuals. In addition, we considered BRIC2 patients (BRIC2: benign recurrent intrahepatic cholestasis type 2) who are mostly asymptomatic but may develop symptoms of cholestasis following medical incidents. For the analysis of drug-induced cholestasis we analysed administration of cyclosporine A (CsA) which is known to cause cholestasis, due to competitive inhibition of canalicular transporters.

## Materials and Methods

### PBPK Modelling

Physiology-based pharmacokinetic (PBPK) models are computational models that mathematically describe the physiological processes underlying absorption, distribution, metabolism, and excretion (ADME) of compounds such as xenobiotics or endogenous molecules within the body of an organism at a large level of physiological detail. The whole-body concept includes all major organs of the modelled organism, such as the liver, heart, or kidney. The organs are further subdivided into plasma, red blood cells, interstitium, and intracellular space. Since PBPK models are knowledge-based most parameters describing the anatomy or physiology of the body such as organ volumes, surface areas, or blood perfusion rates are taken from curated data collections which are provided to the user within PBPK software. The different organs in a PBPK model are interconnected through the vascular circulation. Such a high level of detail of the organism physiology allows a mechanistic representation of complex biological systems and phenomena as well as the individualisation of patient models for example through the consideration of specific phenotypes. Physiologically relevant and tissue-specific active ADME processes such as enzymatic metabolism and transport can also be considered in PBPK models. Tissue-specific gene expression data can be integrated as a surrogate for enzyme and transport protein levels in active processes (Meyer et al. 2012). Besides physiological and anthropometric information of the modelled organism, substance-specific physicochemical parameters like the molecular weight, solubility, or lipophilicity are used as input parameters during PBPK model development. In particular, these values are used to calculate passive diffusion processes across membranes or organ-plasma partitioning coefficients in the distribution models typically underlying PBPK models. (Kuepfer et al. 2016)

The software tools MoBi® and PK-Sim® from the Open Systems Pharmacology (OSP) platform were used for the development of the PBBA model (Open Systems Pharmacology Suite). The latest versions of PK-Sim® and MoBi® are freely available under the GPLv2 License. The basic PBBA model describing mainly passive distribution of GCDCA was established within PK-Sim®, while the additional endogenous processes, i.e. BA synthesis and gall bladder emptying events, were implemented in MoBi®. For the population simulations a virtual population of 1,000 healthy individuals with varied anthropometric properties (Age: 20-60 years, females: 50%, BMI: 19-25kg/m2) and reference concentrations for all transporters was constructed in PK-Sim®. Up to 10% variation was allowed for the transporters abundance. Population simulations and model analyses were performed in Matlab (Version 8.5.1.281278; The MathWorks Inc., Natick, MA).

### Competitive Inhibition of BSEP transport by Cyclosporine A

A PBPK model of Cyclosporine A (CsA) built with PKSim was coupled to the PBBA model to simulate the effects of CsA on BA levels. Additionally, a term describing the competitive inhibition by the drug on BSEP transport kinetics was introduced to the combined model as follows:

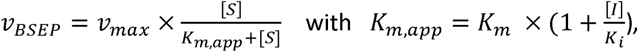

Where [*S*] is the concentration of BSEP substrate GCDCA, [*I*] is the concentration of the inhibitor CsA, *K*_*i*_ is the inhibitor’s dissociation constant, and *K*_*m*_ is the Michaelis Menten constant (Berg et al. 2018)

### Experimental Data

Experimental data from literature describing BA levels in the blood plasma of healthy individuals were used for parameter estimation and model validation. BA plasma levels under fasting conditions were used to identify the basal level of systemic BAs (Bathena et al. 2013).

In addition, various studies that measured postprandial plasma BA profiles of three subsequent meals in healthy male individuals (Hepner and Demers 1977) (Schalm et al. 1978), healthy woman (Angelin and Björkhem 1977), pregnant woman, and diseased volunteers were used to identify the system dynamics of circulating BA level in the human body. Furthermore, we used experimental data of studies not used for parameter identification to validate our model predictions and to additionally assess the variability of individual BA blood plasma levels. (Galeazzi et al. 1980), (Larusso et al.), (Gälman et al. 2005), (Ponz de Leon et al. 1978) and (Salemans, J. M. J. I. et al. 2009). If necessary, experimental plasma BA data were extracted from the original publications with the web-based WebPlotDigitizer tool (version 3.9; Ankit Rohatgi, Austin, TX, USA, freely available under the GPLv3 License http://arohatgi.info/WebPlotDigitizer). Notably, the various studies measured different BA conjugates making it difficult to compare the studies directly. In the study of (Hepner and Demers 1977) glycine conjugates of cholic acid (CA), chenodeoxycholic acid (CDCA), deoxycholic acid (CDA) and sulpholithocholic acid (LCA) were identified, as such representing only a subset of the complete BA pool. Another study (Schalm et al. 1978), investigated postprandial plasma BA profiles in five healthy as well as in pregnant and diseased volunteers and measured chenyl- and cholyl- conjugates. Whereas yet others (Angelin and Björkhem 1977) measured postprandial plasma BA profiles in five healthy women and quantified CA, CDCA and DCA without amidation and sulphation. The measured BA species vary considerably among various studies such that we normalized the postprandial BA profiles to allow a comparison of the different data. Therefore, a percentage scaling factor was calculated from literature for scaling all datasets to the fraction of summed conjugated cholic, chenodeoxycholic, and deoxycholic acid as far as the study description allowed (Table S1, Figure S1) (Bathena et al. 2013). However, it is known that different BA species do not have the same kinetics. Thus, this approach introduces a systemic error leading to a reduced fit quality of the model to the experimental data but is sufficient to describe the overall BA metabolism behaviour.

## Results

### Physiology-based model of bile acid metabolism

We started our analysis by establishing a physiology-based bile acid (PBBA) model at the whole-body scale for a healthy reference individual. The overall workflow of the study in terms of model development and subsequent analyses is presented in Figure 1. A reference model of BA metabolism in healthy humans was developed based on physicochemical properties of a representative BA and known physiological processes. Subsequently, a virtual population of 1,000 individuals was created to assess the variability in post-prandial BA levels. In the next steps, the impact of the BRIC2 and PFIC2 mutations as cause of DILI predisposition and CsA administration on BA levels were predicted and analysed.

**Figure 1.**
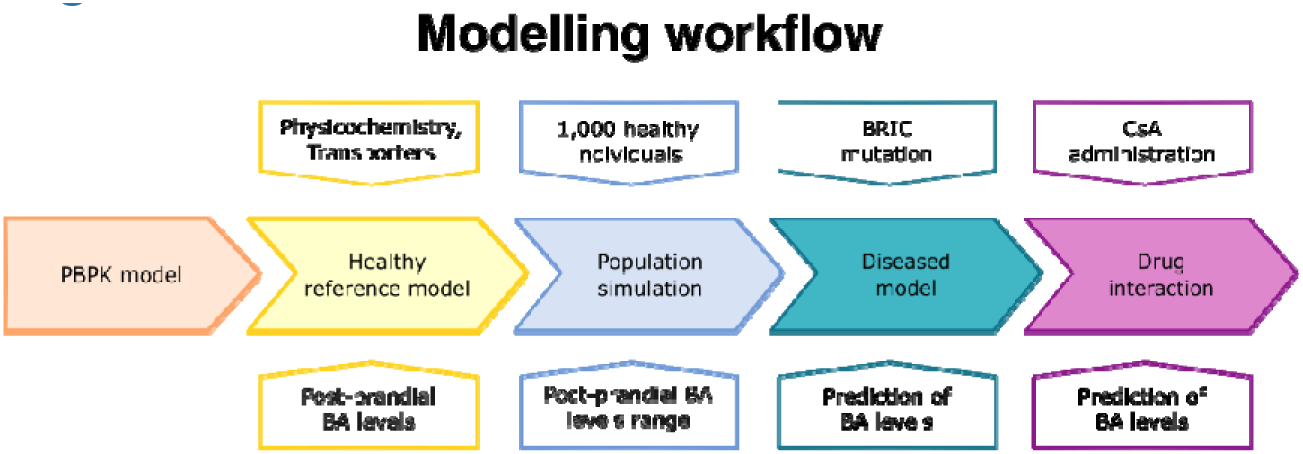
Workflow of the study. The five steps of model development are depicted: basic PBPK model, healthy reference model, population simulation, diseased model for BRIC, and drug interaction with Cyclosporine A. The upper row of boxes depicts the inputs for the different model stages (middle row). The lower row depicts the outputs of the model simulations.

The reference model includes synthesis, circulation and excretion of an exemplary BA. Here, glycochenodeoxycholic acid (GCDCA) was chosen which is the most abundant BA and accounts for 20 % of the total BA pool (Bathena et al. 2013). On one hand, this allowed us to limit the model complexity to the key physiological processes as such improving identifiability of the free model parameters. On the other hand, the consideration of a single surrogate BA species allowed the integration of heterogeneous literature data that analyse different BA species through the earlier described scaling. Following best practice guidelines for PBPK model building, physicochemical parameters of GCDCA like molecular weight, solubility, lipophilicity (logP), and plasma-protein binding (fraction unbound) (Table 1) were used to parametrise the compound properties of the PBPK model for small molecules. Thus, passive transport processes as well as organ-plasma partitioning can be directly calculated using an appropriate distribution model (Kuepfer et al. 2016).

**Table 1.**
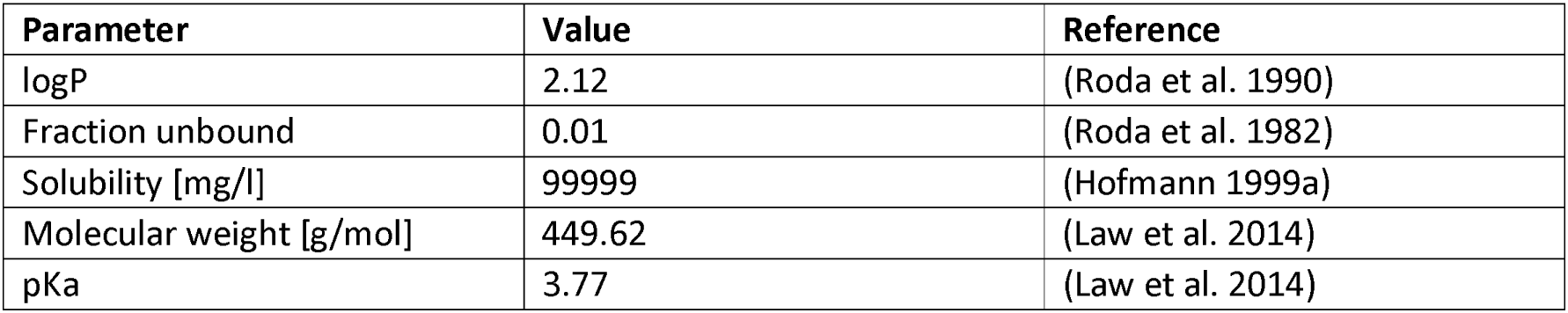
Physico-chemical parameters of bile salt GCDCA

| Parameter | Value | Reference |
| --- | --- | --- |
| logP | 2.12 | (Roda et al. 1990) |
| Fraction unbound | 0.01 | (Roda et al. 1982) |
| Solubility [mg/l] | 99999 | (Hofmann 1999a) |
| Molecular weight [g/mol] | 449.62 | (Law et al. 2014) |
| pKa | 3.77 | (Law et al. 2014) |

BA metabolism is a nearly closed circuit including initial synthesis, transformation, diffusion and multiple active transport processes (Figure 2). Within the body, BAs undergo continuous enterohepatic circulation connecting liver and gastrointestinal tract through the gut-liver axis. Due to very effective recycling, only around 5% of the BA pool is excreted over 24h mainly via faeces (Houten et al. 2006). To compensate for the daily loss of BAs, a continuous synthesis reaction was introduced to the model. This formation of GDCDA is represented by a constant synthesis in the intracellular space of the liver. In vivo, this synthesis rate accounts for cytochrome P450-mediated oxidation of cholesterol and subsequent conjugation with glycine within the liver (Martinot et al. 2017), (Kullak-Ublick et al. 2004). Upon synthesis, BAs are actively secreted in hepatocytes or enterocytes. In hepatocytes, they are either transported to bile canaliculi (apical) or the liver sinusoids (basolateral) while in enterocytes, BAs are secreted to either the gut lumen (apical) or the blood plasma (basolateral). Notably, these transporter-mediated processes are key steps in enterohepatic circulation which have a significant impact on dynamics and mass distribution of the BA pool. These active processes can be also structurally presented in PBPK models (Meyer et al. 2012). In total, four active transport processes were included in the PBBA model: (1) The bile salt excretion pump (BSEP) on the apical membrane of hepatocytes, (2) the Na^+^-taurocholate co-transporting polypeptide (NTCP) on the basolateral membrane of hepatocytes, (3) the human ileal the bile acid transporter (ASBT) apically in the ileum mucosa, and (4) the organic solute and steroid transporter (OSTα/β) basolaterally in the ileum mucosa (Kullak-Ublick et al. 2004; Rao et al. 2008). A fraction of 65% of biliary excreted BAs was assumed to be collected in the gall bladder while the remaining fraction is directly transported to the duodenum (Hofmann 1999a). Gallbladder emptying is triggered by meal ingestions. In all simulations, three meals over 24h representing breakfast, lunch and dinner were considered. To close the overall mass balance, faecal and renal excretion of GCDCA were implemented in the model by passive transport and active linear clearance, respectively. (Table 2) Altogether, the initial PBPK model of GCDCA structurally describes continuous BA synthesis as well as enterohepatic circulation through the liver and the GI tract including re-absorption from the ileum. It should be noted that as such the model is similar to a typical PBPK model for xenobiotic drugs even though distribution and excretion of an endogenous compound are considered here. This similarity is a particular advantage of our approach since the basic PBPK model already includes a detailed physiological representation of the gastro-intestinal tract involving several segments to quantitatively describe dissolution of tablets in the gastro-intestinal tract (Thelen et al. 2011). This is of outmost importance to physiologically describe re-absorption of BAs from the gut lumen during enterohepatic circulation.

**Figure 2.**
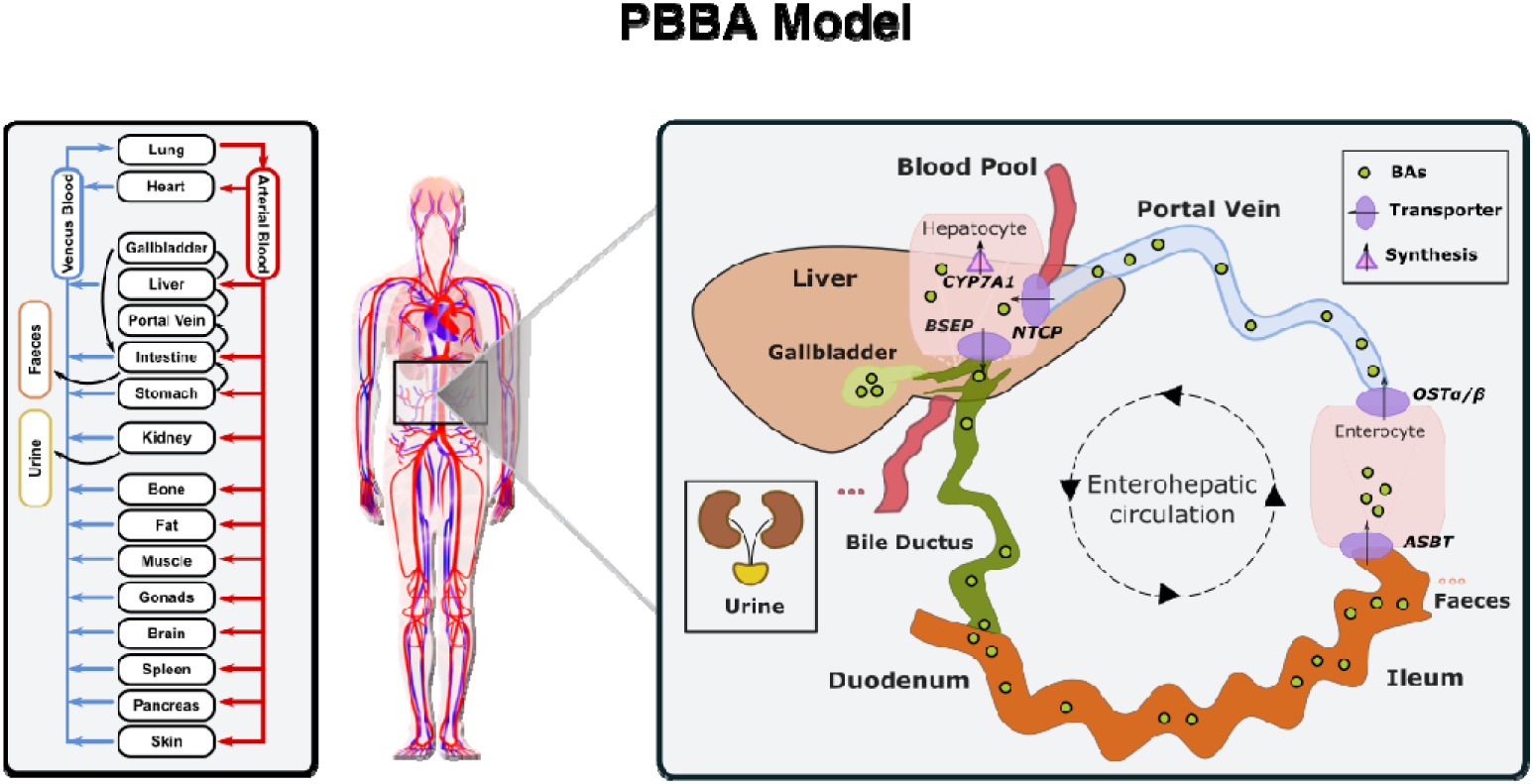
Whole-body physiology-based pharmacokinetic model of the bile acid metabolism. The platform includes a PBPK model of the bile acid GCDCA, with biosynthesis via CYP7A1 in the liver, active transport processes via BSEP, ASBT, OSTα/β, and NTCP, faecal and renal excretion. GCDCA is stored in gallbladder and partially secreted directly into the duodenum and is reabsorbed along the intestine (enterohepatic circulation). Emptying of gallbladder is triggered by food intake.

**Table 2.** PBBA model parameters.

| Parameter | Value | Start value |
| --- | --- | --- |
| $K_m$ (BSEP) [$\mu\text{mol/l}$] | 5 | 4 (Kullak-Ublick et al. 2004) |
| $K_m$ (ASBT) [$\mu\text{mol/l}$] | 0.5 | 50 |
| $K_m$ (NTCP) [$\mu\text{mol/l}$] | 1 | 6 (Kullak-Ublick et al. 2004) |
| $K_m$ (OST$\alpha/\beta$) [$\mu\text{mol/l}$] | 7.5 | 50 |
| $k_{cat}$ (BSEP) | 300 | 100 |
| $k_{cat}$ (ASBT) | 5 | 100 |
| $k_{cat}$ (NTCP) | 125 | 100 |
| $k_{cat}$ (OST$\alpha/\beta$) | 9000 | 1000 |
| Body weight [kg] | 73 | Not fitted |
| Age [years] | 30 | Not fitted |
| Height [m] | 1.76 | Not fitted |
| GB Volume [l] | 0.02 | Fixed (van Erpecum et al. 1992) |
| Refilling time [min] | 147.8 | 419 |
| Emptying half life [min] | 69.98 | 69.98 |
| Synthesis rate [$\mu\text{mol/min}$] | 0.78 | Fixed (Kullak-Ublick et al. 2004) |
| Renal excretion [$\mu\text{mol/l/min}$] | 981.30 | 100 (Bathena et al. 2015) |
| $K_i$ (CsA) [$\mu\text{mol/l}$] | 2 | 2 (Böhme et al. 1993) |
| Distribution model | PKSim standard |  |

Next, uninformed model parameters were identified in order to accurately describe the dynamics of the BA metabolism in a healthy reference individual. Importantly, only a limited set of model parameters had to be considered since the model relies on large datasets of physiological and physicochemical information (Table 2). Measurements of basal and postprandial BA concentration levels from literature were used to evaluate the agreement of the computational simulations with the scaled experimental data (Hepner and Demers 1977; Ponz de Leon et al. 1978; Schalm et al. 1978). The comparison between the simulated kinetics of BA levels over 24 hours with reported data is shown Figure 3. Experimental data of post-prandial BA profiles from two studies are shown in which healthy individuals were fasted overnight and given three meals at 8:00, 12:00, and 16:00 h. Following parameter identification, the model could describe the plasma bile acid dynamics well despite a significant level of variability in the experimental data. The peak concentrations are met with sufficient accuracy as well as the corresponding trough levels between the meals and the dynamics of gallbladder emptyings.

**Figure 3.**
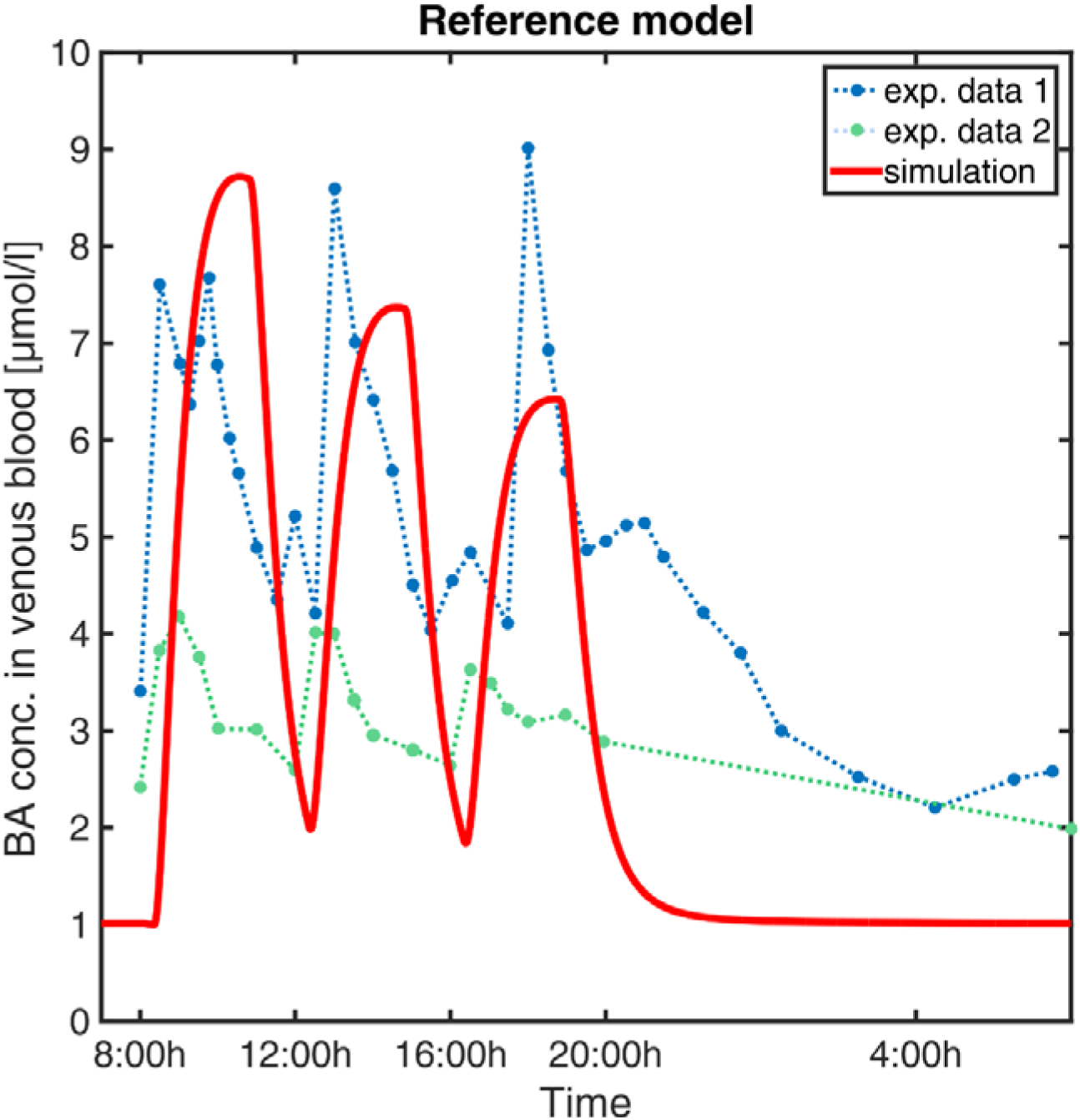
Simulation of venous blood plasma BA levels in a human reference individual. The PBBA model was simulated with three meals per day given at 8, 12, and 16 o’clock and simulated BA concentrations in venous plasma (red solid curve) are compared with reported values from (Hepner and Demers 1977) (exp. data 1, dark blue points connected by dashed line) and (Ponz de Leon et al. 1978) (exp. data 2, green points connected by dashed lines).

We next verified whether several physiological reference measurements of the BA metabolism such as total BA pool size, cycling times and concentrations in various compartments could be described with the model. A series of clinical parameters was hence collected from the literature and used for fitting and comparing to corresponding values calculated from the simulation results (Table 3). Even though the model is an open system with a complex dynamic behaviour a good agreement was achieved. This accordance of physiological reference values between the literature values and the simulation outcomes is a very strong indication for the correctness of the overall model in terms of both mass balance and dynamics (Figure 3).

**Table 3.** Physiological reference measurements. Values with (*) were converted to model units.

| Parameter | Literature | Model | Reference |
| --- | --- | --- | --- |
| BA conc. in venous blood [$\mu\text{mol/l}$] | [0.9, 8.4] | [1.68, 8.91] | Var. Sources (see M&M) |
| BA conc. in portal vein blood [$\mu\text{mol/l}$] | [2.8, 33.2] | [3.95, 34.5] | (Angelin et al. 1982) |
| Faecal excretion rate [$\mu\text{mol/min}$] | 0.72* | 0.72 | (Dancygier 2003) |
| BA pool size [$\mu\text{mol}$] | [4250, 6672]* | 5697.69 | (Beuers et al. 1992;<br>Bisschop et al. 2004) |
| Avg. secretion rate per meal [mmol/h] | 5 | $\sim 1.2$ | (Hofmann 1999a) |
| BA conc in gallbladder [$\mu\text{mol/l}$] | [3000; 100,000] | [25;2800;5000] | (Hofmann 1999a; Jansen<br>et al. 2017) |
| BA conc. in intestinal lumen [$\mu\text{mol/l}$] | [2000, 10,000] | [75, 5175.02] | (Hamilton et al. 2016;<br>Hofmann 1999a) |
| BA conc. in liver cells [$\mu\text{mol/l}$] | 1-2; < 3 | [0.23, 1.07] | (Hofmann 1999a, 1999b) |

### Population Simulation of the PBBA model

At this stage, the PBBA model describes BA profiles in an average adult individual. To also account for the high inter-individual variability within the clinical data and to test the robustness of the PBBA model, a population simulation was additionally performed. Figure 4 compares the BA levels per meal from the population simulation to the experimental data from five studies (see Materials & Methods). To this end, the measurements of both the experimental data from literature and the simulation results were assigned to a first, second, and, third meal whenever possible. Single BA measurement points in plasma (one symbol per study) as well as boxes indicating the 25^th^ and 75^th^ percentiles for the experimental data (orange) and the simulations (black) are plotted. The median BA concentration per each meal decreases over daytime both in the experimental data and the simulations. The predicted population variability was in a to-be-expected physiological range and matched the experimentally measured BA values.

**Figure 4.**
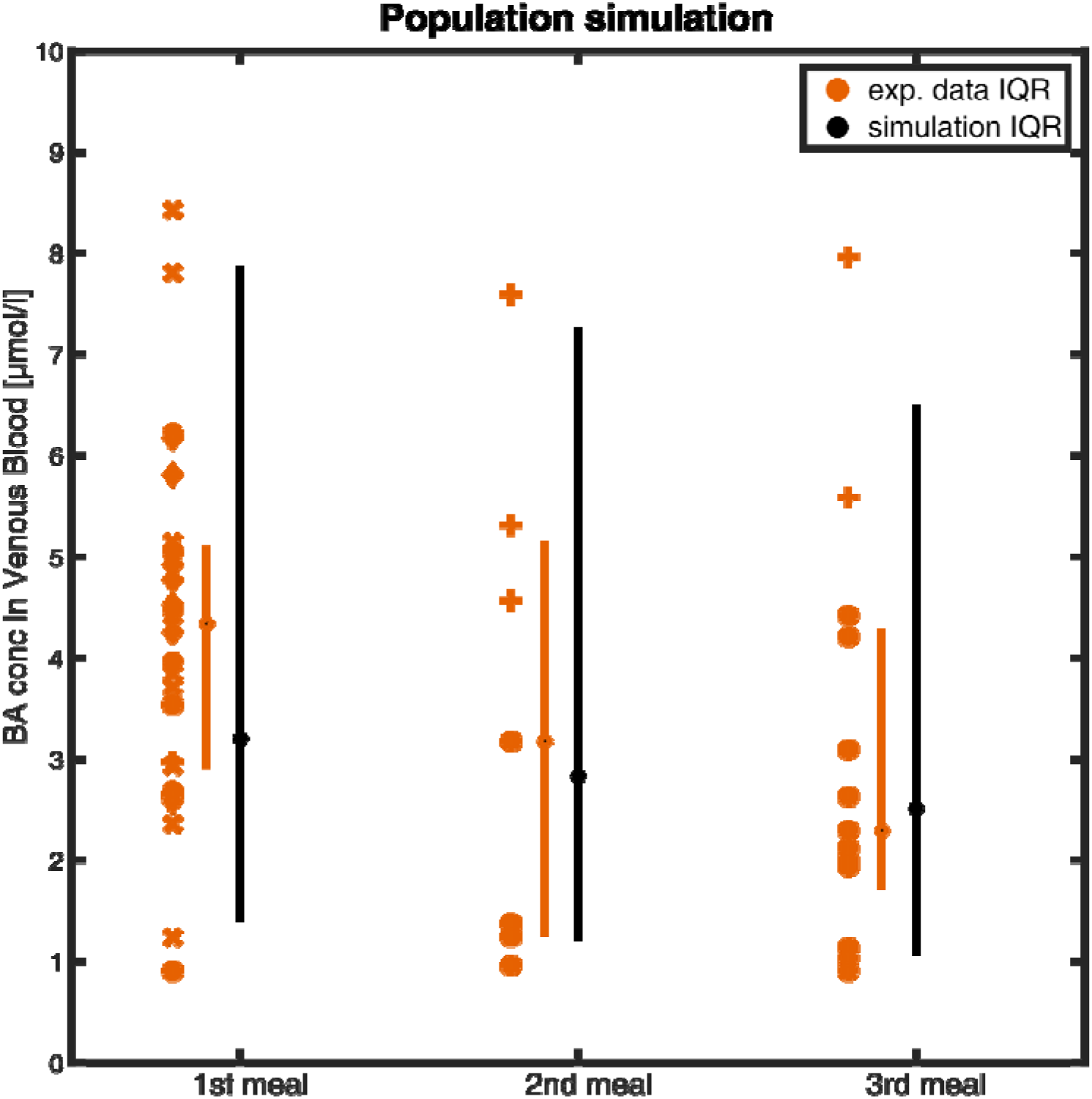
Simulation of venous blood plasma BA levels in a virtual healthy population of 1,000 individuals. BA levels were assigned to the meal after which they were measured or simulated. Symbols represent BA plasma measurements from experimental studies (o = (Angelin and Björkhem 1977), + = (Galeazzi et al. 1980), * = (Gälman et al. 2005), x = (Salemans, J. M. J. I. et al. 2009), ⍰ = (Ponz de Leon et al. 1978)). Boxes represent the interquartiles range of the experimental data (orange) and the simulation (black) with the median marked as dot on the box.

The comparison of the PBBA model with various physiological reference measurements (Table 3) as well as clinical data sets (Figure 4) is a strong indication for the overall correctness of the model for healthy reference individuals. This careful validation hence generates confidence in the further model predictions and investigations.

### Diseased model of Benign Recurrent Intrahepatic Cholestasis type 2

A variety of inborn forms of cholestasis exist in humans (Pauli-Magnus et al. 2005). Depending on the affected protein and the locus of mutation, different types and severity of cholestasis can be present. It has been known for long that carriers of progressive familial intrahepatic cholestasis type 2 (PFIC2) or benign recurrent intrahepatic cholestasis type 2 (BRIC2) have a higher risk of encountering cholestasis as a consequence of other diseases or drug therapies (Srivastava 2014). Both the severe PFIC2 and the milder BRIC2 are caused by polymorphisms of the BSEP-coding gene. The mutations lead to an impaired function of the protein. As a result, PFIC 2 patients usually experience an early onset of cholestasis in their lifetime and often need early liver transplantations. The BRIC2 mutations are usually less severe such that a basal functionality of BSEP remains. However, affected patients have episodes of cholestatic symptoms in their lifetime and can have slightly elevated basal BA plasma levels (Ermis et al. 2010; Hayashi et al. 2016; Zellos et al. 2012).

Based on the PBBA model for a healthy reference individual, we simulated the effect of PFIC2 and BRIC2 on BA levels in different tissues by decreasing the transporter activity in this genotype subgroup. For BRIC2 patients, we reduced the BSEP from 100% to 20-13% of the original BSEP transporter activity to account for the remaining functionality. For the PFIC2-genotype we further reduced the transporter activity of BSEP to 5% (Noe et al. 2005). The simulation results show the relative differences of BA amount in various enterohepatic compartments after simulating the gradual loss of BSEP function (Figure 5). While the downstream compartments of the liver like gallbladder, intestinal tract (not shown), and faeces contain less BAs, the upstream compartments portal vein, venous blood, and urine contain higher amounts of BAs compared to simulation results with 100% BSEP function. The range of BSEP function in BRIC2 individuals is indicated in blue and simulations show that individuals have up to doubled BA levels in blood and up to six-fold increase in the liver cells (Figure 5). It should be noted that it is technically impossible to actually measure BA concentrations in these tissues in BRIC2 patients in clinical practice.

**Figure 5.**
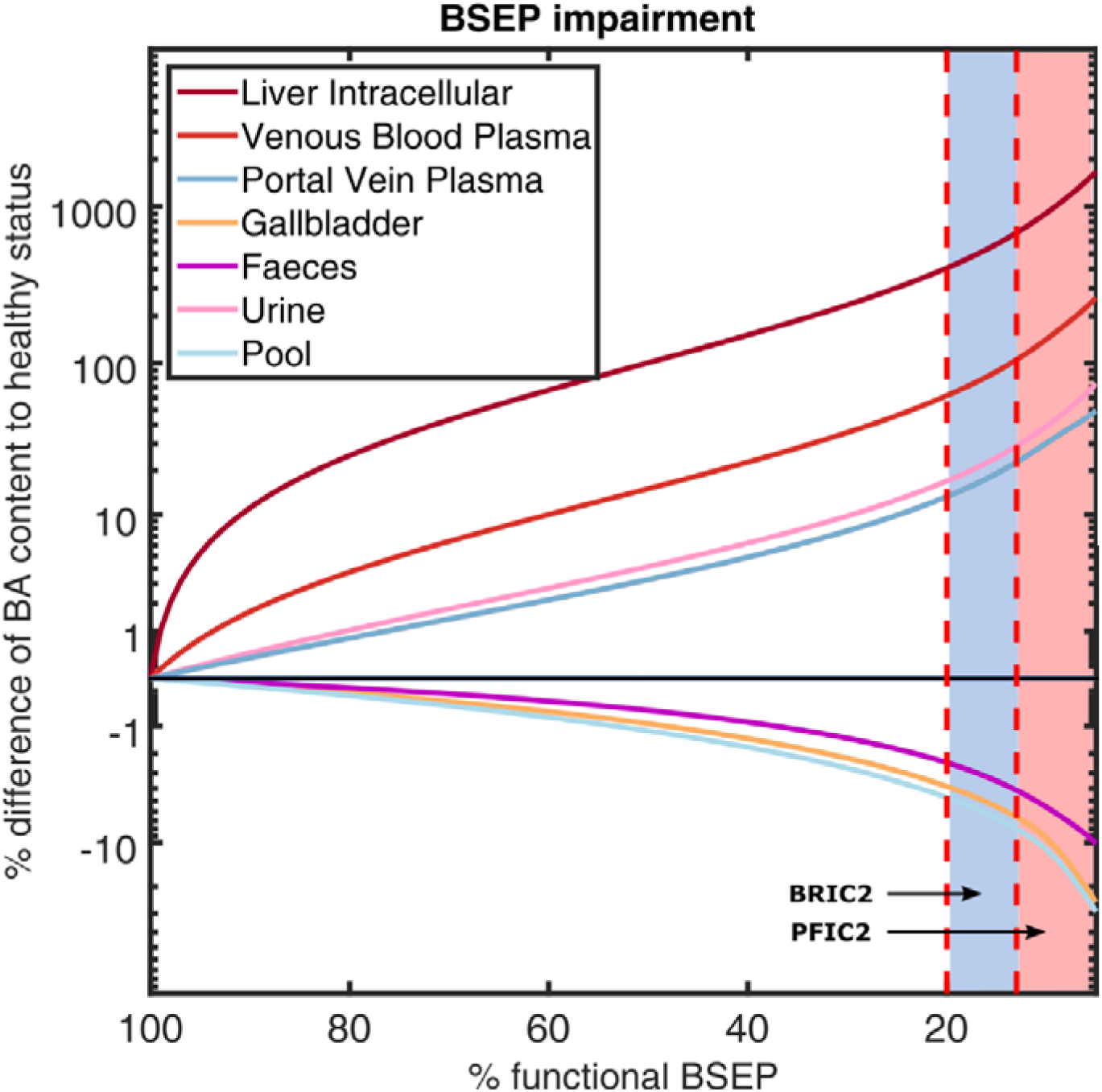
Simulation of BA levels in various compartments with decreasing BSEP function. The mean difference (in % of reference) of BA content in various compartments as well as the total BA pool are plotted over decreasing BSEP function. The reported ranges of BSEP functionality in BRIC2 patients with 20-13% and in PFIC2 with <13% are marked in blue and red, respectively. BA content in faeces, gallbladder, and the total pool decrease with decreasing BSEP function, while BA in liver cells, venous blood, portal vein blood, and urine increase.

### Drug interaction with cyclosporine A

As a further clinical case we considered the influence of cyclosporine A (CsA) administration on BA levels. CsA is known to induce DILIs with different degrees of severity by influencing gene expression of liver enzymes and transporters but also inhibiting transport processes of BSEP competitively. For the investigation of CsA’s impact on BA levels, we coupled the PBBA model with a published PBPK model of CsA (Thiel et al. 2017a). Such interaction models are a frequent application in PBPK modelling (Thiel et al. 2017b), however, it should be noted that in the present case the coupled model simultaneously describes the disposition of endogenous BA species as well as the PK of an exogenous drug. The PBPK model for CsA has been carefully validated before with different PK data for intravenous and oral administration. (Thiel et al. 2017a) (Figure 6). The inhibition of CsA on BSEP was integrated by the introduction of a competitive inhibition term for BSEP kinetics. In addition, the combination of BSEP inhibition by CsA in individuals with a BRIC2-genotype was considered. Figure 7A shows the CsA levels after a bi-daily intravenous dose of 2 mg/kg CsA in venous blood and liver cells. The simulations (Figure 7B) show mildly elevated BA levels in venous and portal vein blood in healthy individuals. A more pronounced effect is seen in the liver cell BA levels with an increase of 22% compared to the untreated case. After CsA administration in BRIC2 patients, BA levels rose up to 739% relative to healthy reference individual if only an additive effect was assumed (Figure 7B). These results suggest that even routine medical incidents may lift BA plasma concentrations in BRIC2 to levels which are no longer tolerable and potentially cholestatic.

**Figure 6.**
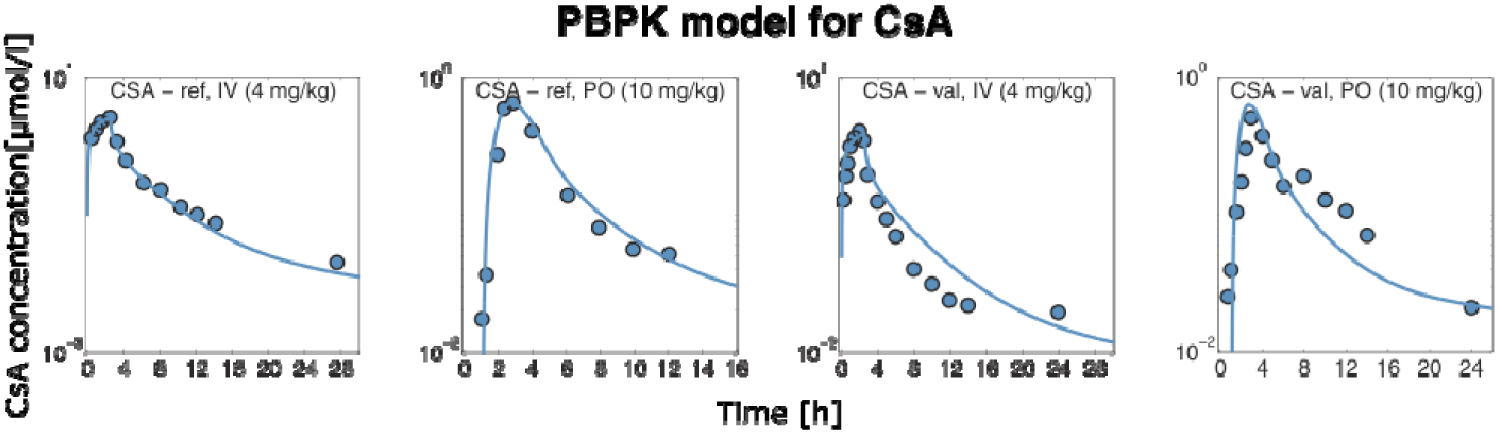
Simulation of cyclosporine A levels in venous blood plasma after intravenous and oral administration of 4mg/kg and 10 mg/kg. Reference datasets used for fitting and validation datasets are shown.

**Figure 7.**
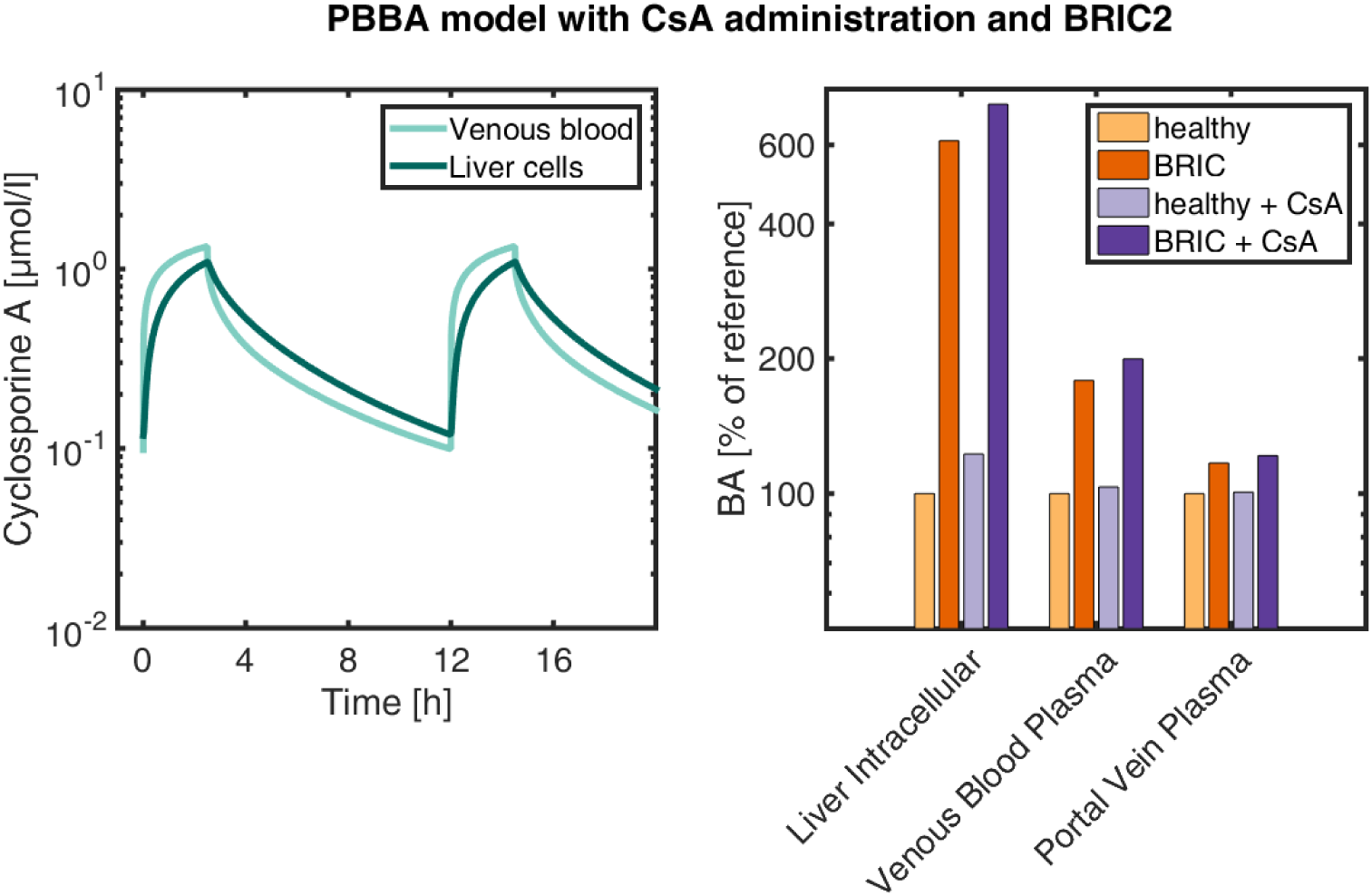
Simulation of the coupled CsA PBPK and the PBBA model models. Right panel: CsA concentrations in venous blood plasma and liver Intracellular space after 2mg/kg iv administration twice daily. Left panel: simulated mean BA (in % of healthy reference) of the reference model, the BRIC2 model and the CsA-coupled models in liver intracellular space, venous blood plasma, and portal vein plasma.

## Discussion and conclusions

Distribution and accumulation of BAs in blood plasma and different tissues are a direct clinical marker for cholestasis. However, targeted sampling at different locations of interest throughout the body is impossible due to ethical and technical restrictions such that a comprehensive picture of BA metabolism is hard to achieve. This is in particular since enterohepatic circulation of BAs is a systemic process which involves multiple consecutive and fine-tuned steps in different organs. Therefore, the causes of aberrant states of BA metabolism are hard to understand in individual patients, which significantly limits the usage of BAs measurement for diagnostic profiling. A computational model that quantitatively describes BA concentration in body fluids at different locations and tissues of the body is hence a valuable tool to generate a comprehensive understanding of the contribution of the various physiological processes involved in BA metabolism. Moreover, such a mechanistic model would be helpful to suggest novel markers of cholestasis in clinical practice. Thus, it would be possible to understand the mechanisms underlying cholestasis, to prevent intoxication and health damage in patients and to design alternative treatment approaches once first DILI symptoms have developed.

We here established a whole-body PBBA model for humans, which allows the estimation of BA exposure in blood plasma and different tissues. The main processes of BA metabolism such as synthesis, excretion, and enterohepatic circulation are mechanistically included in the model. Likewise, as the model builds on PBPK modelling, organ-plasma partitioning is explicitly simulated for different tissues throughout the body. Thus, intracellular concentrations, e.g. in the liver where accumulation occurs and the chances of damage are high are directly available. Additionally, concentrations in other potentially vulnerable organs such as skin may also be simulated. In this way, the risk of complications like for example pruritus can be evaluated. We parametrised the model based on a comprehensive set of publicly available plasma BA data and validated it afterwards with an independent data set not used in model establishment. We showed, that our model is able to reproduce the dynamic postprandial BA levels in healthy individuals, as well as to predict BA levels for different clinical cases such as different genotype subgroups and for drug-administration. To this end, genotype-specific functionality of BSEP transporter was simulated and confirmed the predisposition of these subgroup towards drug-induced cholestasis by elevated BA levels in blood and in liver cells. The additional administration of CsA to this patient group increased the BA levels even further and with this the cholestasis risk. Moreover, it is only through the computational model that BA exposure at specific sites of the body can be quantified through the simulations. The computational model allowed a systematic consideration of different degrees of impaired BSEP activity in BRIC2 and PFIC2, which could otherwise not have been analysed at such a high step density. In addition, the here developed physiology-based computational model is not data-driven but knowledge based since a lot of prior physiological information is included in the models during its development. For this reason, it is also possible to hypothetically consider specific questions like functional changes or alterations in environmental conditions which have not explicitly considered during model establishment itself.

The first attempts to mathematically model the BA metabolism were published in the early 1980s (Cravetto et al. 1988; Hofmann et al. 1983). These models were detailed in the description of the BA species and the enterohepatic circulation but lacked mechanistic knowledge regarding the whole-body physiology and relevant transporters. Recent models make use of a simplified representation of the body physiology and do not include organs or their sub-compartments such as intracellular or interstitial space. (Sips et al. 2018; Woodhead et al. 2014). Moreover, such models are mostly data-driven instead of knowledge-based which limits translation to new indications, patient subgroups or clinical scenarios such as BRIC2 and PFIC2 patients which were considered in this study. Since our focus lies on the basic physiological mechanisms of cholestasis development, none of the general yet secondary clinical biomarkers like ALP have been considered in our model (Longo et al. 2016). Instead, we aimed towards a description of the actual defect and not the indirect consequences of the tissue damage induced by accumulated BA. In contrast to data driven approaches (Sips et al. 2018), the presented PBBA model is knowledge-based and relies on huge prior information what makes the identified parameters of the model reliable.

In the future, the PBBA model will also help to explain the causalities in idiosyncratic cases of DILI such as genetic or physiological predisposition of individual patients. Functional consequences of alterations such as different genotypes or diseases can be mechanistically represented in physiology-based modelling (Lippert et al., 2012) allowing for example to describe cases of DILI well beyond intrinsic, i.e. predictable dose-related DILI. Since PBPK models allow the inclusion of patient specific physiological information the PBBA model is very well suited to analyse cases of idiosyncratic drug-induced cholestasis. In particular, the model allows to describe individual drug exposure in off-target tissue as a consequence of a patient’s anatomy, physiology, lifestyle, gender or age. In addition to the listed advantages, the used PBPK framework can be in general translated from the current human PBBA model to other species like mice to support model-based experimental design (Thiel et al., 2015).

However, the current version has a number of shortcomings. On one hand, the population simulation showed that the simulated variability is higher than the measured one. That could be overcome by adapting the synthesis rate to changing liver volumes instead of fixing it. On the other hand, basolateral transport processes for BA excretion from the hepatocytes was neglected since it is difficult to differentiate because only a net transport rate into the hepatocytes can be reliably identified. For this reason, counteracting processes which could potentially reduce the effect of BSEP functional impairment in BRIC and PFIC2 patients have not been considered so far (Figure 5). In this version of the PBBA, we only considered GCDCA as a surrogate BA. However, it is known that different BA species do not have the same kinetics. Therefore, this approach may introduce a systemic error leading to a reduced fit quality of the model to the experimental data. Still, it should be noted that the model is sufficient to describe the overall behaviour of BA metabolism. It can also be argued that, the smaller peaks secondary to the main meal peaks (Figure 3) are not reflected by the model. This is probably due to the not included BA species which show different dynamics. Therefore, future modifications of the model will include differentiation in various BA metabolites. This would be of particular interest since BAs are continuously transformed and accumulate differentially in various tissues all over the body. The prediction of such shifts in BA composition in specific tissues like the liver based on simpler blood samples could be a useful biomarker in cholestasis assessment. For such a differentiated BA pool the necessary metabolisation steps which are catalysed by the intestinal microbiome need to be integrated. However, the tools for the vertical integration of metabolic network models into PBPK models already exist and can be added to the current PBBA model straight forward (Krauss et al., 2012; Cordes et al., 2018). Further modifications of the presented PBBA model could include circadian gallbladder emptyings overnight which have been neglected in the present version of the model. Therefore, the simulated nightly BA profiles are not as reliable as of now. Additionally, different meal compositions could be considered to specifically trigger different responses. This could also include the effect of change in lifestyle on the composition of the intestinal microbiome and subsequent changes in BA composition (Wahlström et al. 2017). Furthermore, specific preclinical in vitro data can be integrated and used for in vitro-in vivo translation of omics data (Kuepfer et al., 2017). This will ideally involve time series of omics data which could be contextualized in the model to track pathogeneses. The clinical cases shown in this work, however, show that the current model can already be applied to analyses of clinical relevance. In particular the model may be used to support the early identification of potential cholestatic drug effects together with targeted experimental data. We therefore think that the presented model provides an important platform for model-based analyses of BA metabolism in the future.

